# Genomic hypervariability of phage Andromeda is unique among known dsDNA viruses

**DOI:** 10.1101/619015

**Authors:** Damian J. Magill, Leonid A. Kulakov, Timofey A. Skvortsov

## Abstract

A new lytic bacteriophage Andromeda, specific to the economically important plant pathogen *Pseudomonas syringae*, was isolated and characterised. It belongs to the *Podoviridae* family, *Autographivirinae* subfamily and possesses a linear dsDNA genome of 40,008 bp with four localised nicks. Crucially, Andromeda’s genome has no less than 80 hypervariable sites (SNPs), which show genome wide distribution resulting in heterogenous populations of this phage reminiscent of those of RNA virus quasispecies. Andromeda has no nucleotide sequence homology to phage phiNFS, a member of *phiKMVviruses*, in which a similar phenomenon was discovered. We show that Andromeda and Andromeda-related phages form a group within the *Autographivirinae*, designated here as the “ExophiKMVviruses”. The “ExophiKMVviruses” were revealed to share conservation of gene order with core *phiKMVviruses* despite their sequence-based relationship to SP6-related phages. Our findings suggest that genomic hypervariability might be a feature that occurs among various *Autographivirinae* groups.

## Introduction

The *phiKMVviruses* constitute a ubiquitously distributed group of bacteriophages closely related to *Teseptimaviruses* and *Zinderviruses*. These three large groups and a number of smaller ones fall under the *Autographivirinae* subfamily, the major difference between them being the localisation of their single subunit RNA polymerase enzymes. With respect to *phiKMVviruses* in particular, the majority of phages infect *Pseudomonas* strains, with first representatives being described in 1980s [1]. *phiKMVviruses* are of interest for various reasons. Those infecting *Pseudomonas aeruginosa* exhibit a number of interesting features including the presence of four localised single-strand interruptions (nicks) associated with the consensus sequence 5’-CGACNNNNNCCTACTCCGG-3’ [1, 2]. The more recent study revealed that at least some phiKMV phages exhibit hypervariability associated with specific genomic regions [3]. These hypervariable loci were found in genes associated with phage adsorption/host specificity and may therefore influence the evolutionary pathway of the phages [3].

This study describes the biological properties of a novel phage, Andromeda, which infects *Pseudomonas syringae*. The presence of a large number of hypervariable sites in the Andromeda’s genome and its relationship to *phiKMVviruses* are investigated.

## Materials and Methods

### Phage Isolation and Purification

Bacteriophage Andromeda was isolated by enrichment of 100 ml of water from River Lagan (Belfast, Northern Ireland) in cultures of *Pseudomonas syringae* pv *syringae* Van Hall followed by plating using the double agar overlay plaque assay [4]. Pure isolates of the phage were obtained by conducting three rounds of phage propagation from individual plaques. To obtain high titre preparations of the phage, cultures of *P. syringae* (OD_600_ = 0.3) were inoculated at a MOI = 0.01. Chloroform was added immediately after culture lysis (∼4 hours) to maximise yield. Resistance of the phage particles to chloroform treatment was determined in separate experiments. The phage lysate was concentrated by PEG precipitation, and purified using CsCl density gradient centrifugation as described by Sambrook and Russell [4]. Bands were extracted and dialysis ensued with modified SM buffer (100 mM NaCl, 50 mM Tris, 10 mM, 8 mM MgSO_4_, 10 mM CaCl_2_) using Amicon Ultra-15 100kDa filters (Millipore). Phage concentrates were treated with SDS and proteinase K, after which phenol-chloroform DNA extraction followed by ethanol precipitation was used to obtain phage DNA [4]. DNA was further purified twice prior to sequencing using the MoBio DNA Purification Kit according to the manufacturer’s instructions.

### Electron Microscopy

Carbon-coated formovar grids were subjected to hydrophilic treatment with poly-L-lysine for 5 min. Upon drying, CsCl purified and dialysed phage suspensions were deposited on the grids and successively stained with 2% uranyl acetate (pH 4.5). Samples were observed using a Phillips CM100 transmission electron microscope at 100 KeV.

### Determination of Adsorption Coefficient

Determination of adsorption efficiency of phages was carried out using the method described in [5]. All adsorption assays were conducted in triplicate with final adsorption constant calculated from the mean of the replicate constants.

### One-step Growth Curves

Determination of bacteriophage one-step growth curves was conducted according to the established protocol. Bacterial cultures were grown until OD_600_ = 0.3 (10^8^ CFU/ml) and inoculated with phage to give MOI ≥ 1. Adsorption was allowed for 5 min at 37°C followed by incubation with aeration at the same temperature. Duplicate 100 μl samples were taken every 5 minutes for 90 minutes. One of each samples were treated with 150 μl of chloroform and both were plated on the lawns of *P. syringae* using the double agar overlay plaque assay. Burst size was calculated by dividing the average of the values following the growth phase of the curve (the step) by the average of the baseline values before the first increase was observed in titre. The average of triplicates was then taken as the burst size. Latent periods were taken as the point in the graph of the untreated phage extraction whereby a rise in titre was observed, whereas the eclipse period was determined from the chloroform treated curve.

### Visualisation of localised single-strand interruptions

Detection of localised nicks were conducted as described earlier [2]. Phage DNA (3 μg) underwent overnight ligation at 4°C in a final volume of 20 μl. NaOH was added to reactions to a final concentration of 50 mM. This was also done for 1.8 μg of unligated DNA. Both samples were incubated for 5 min on ice and immediately analysed by gel electrophoresis (0.9% agarose).

### Library Preparation and Sequencing

Sequencing libraries were prepared from 50 ng of phage genomic DNA using the Nextera DNA Sample Preparation Kit (Illumina, USA) at the University of Cambridge Sequencing Facility. A 1% PhiX v3 library spike-in was used as a quality control for cluster generation and sequencing. Sequencing of the resulting library was carried out from both ends (2×300 bp) with the 600-cycle MiSeq Reagent Kit v3 on MiSeq (Illumina, USA) and the adapters trimmed from the resulting reads at the facility.

### Sequence Assembly and Bioinformatics

Sequencing reads underwent initial quality checking using FastQC (https://www.bioinformatics.babraham.ac.uk/projects/fastqc/) followed by stringent trimming parameters. Reads with a Q-score < 20, containing N’s, and with a final length < 40bp were discarded using Trimmomatic v0.32 [6]. Assembly was carried out using Geneious R8 (Biomatters, New Zealand) and checked for potential misassembly by comparison with output from SPAdes v3.7.0 [7]. Genome length contigs were input to the scaffolding tool SSPACE to extend these in order to ensure the resolution of genome ends [8]. In addition, the sequence obtained was utilised as a reference for read mapping using bbmap v35.x (https://jgi.doe.gov/data-and-tools/bbtools/) and subsequent manual inspection in IGV to check for potentially ambiguous regions [9]. ORFs were classified using the intrinsic Artemis [10] ORF classification and GeneMark.hmm [11] followed by manual inspection, ORF borders correction and verification of ribosomal binding sites. Gene annotation was carried out using BLASTp, Delta-BLAST, HMMER, InterProScan and HHpred [12–15]. tRNAs were classified using tRNA-scanSE and Aragorn [16, 17], and rho-independent terminators were identified using Arnold [18] with an energy cutoff of -10 employed, following by manual inspection of the output from subsequent Mfold analysis and adjustment as necessary [19]. To identify putative regulatory sequences, 100 bp upstream of every ORF was extracted using an in-house script and analysed using both neural network promoter prediction for prokaryotic sigma 70 promoters and Multiple EM for Motif Elicitation to identify phage specific promoters [20]. Phage Andromeda assembled genomic sequence was deposited to NCBI Genbank (Genbank accession number: KX458241; RefSeq accession number: NC_031014). A reannotated genome of phage Andromeda in Genbank format can be found in Supplementary File 1.

### Phylogenetic analysis

In order to taxonomically annotate the newly isolated phage and determine its phylogenetic relationships with other bacteriopahges, two conserved Andromeda proteins, the single subunit RNA polymerase (RNAP, YP_009279548.1) and the terminase large subunit (TerL, YP_009279565.1), were used to generate phylogenetic trees. BLASTp search with each of these proteins against *Podoviridae* available in RefSeq at the moment of writing was conducted and only phage genomes containing copies of both TerL and RNAP with high enough similarity (e-value ≤ 1e-5; query coverage ≥ 75%; percent identity ≥ 25%) to corresponding Andromeda’s proteins were retained for subsequent analyses; 147 complete phage genomes were selected in total (Supplementary Table 1). Alignments of the amino acid sequences of RNAP and TerL were carried out in MAFFT using L-INS-I settings [21], TrimAl was used to for automated alignment trimming [22]. The inference of phylogenies was then carried out using IQ-Tree [23], which was allowed to compute the optimal protein model to be used. Visualisation of the resulting tree file was subsequently performed with the Interactive tree of life (IToL) [24]. The GGgenes package (https://CRAN.R-project.org/package=gggenes) for R programming environment (http://www.R-project.org/) was used to generate a series of genome maps.

### Molecular Modelling

Molecular models were constructed from corresponding amino acid sequences using the I-Tasser package [25] and visualised with Chimera [26]. Fidelity of models was assessed using the validation tools of the Whatif server (https://swift.cmbi.umcn.nl/whatif/) and checked for stereochemical clashes within Chimera. Ramachandran analysis was subsequently carried out as an additional point of quality assessment. Superimposition and quantitative similarity of protein structures was provided by the TM-Align algorithm [27].

## Results and Discussion

### Isolation and Biological Characteristics

Bacteriophage Andromeda was isolated from the River Lagan as described in Materials and Methods. Upon infection of *Pseudomonas syringae*, phage Andromeda yields large, clear plaques of ∼3 mm in size. Electron microscopic analysis of Andromeda revealed a virion morphology characteristic of *Podoviridae* (Fig. 1). Specifically, Andromeda was found to possess a capsid of icosahedral symmetry with a length of 61 nm (±1.5 nm) and width of 62 nm (±1 nm) and a detectable tail structure.

**Fig. 1.**
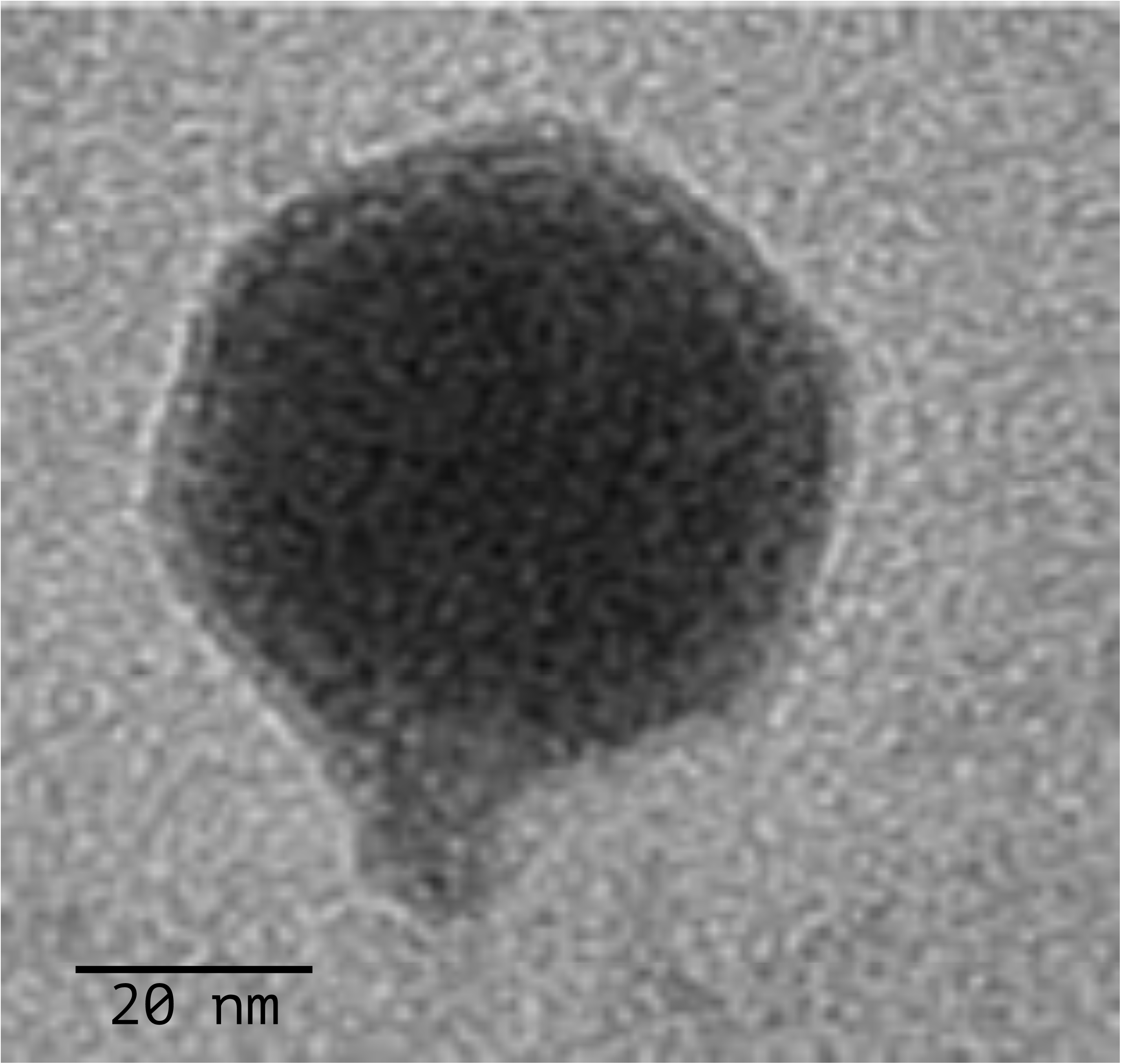
Electron microscopic image of bacteriophage Andromeda.

Analysis of the growth characteristics revealed an adsorption coefficient of 1.48 × 10^−9^ ml min^−1^ (Fig. 2a), eclipse period of 20 – 25 min, and latent period of 25 – 30 min (Fig. 2b). Upon completion of its infection cycle, Andromeda yields on average 70 particles per cell.

**Fig. 2.**
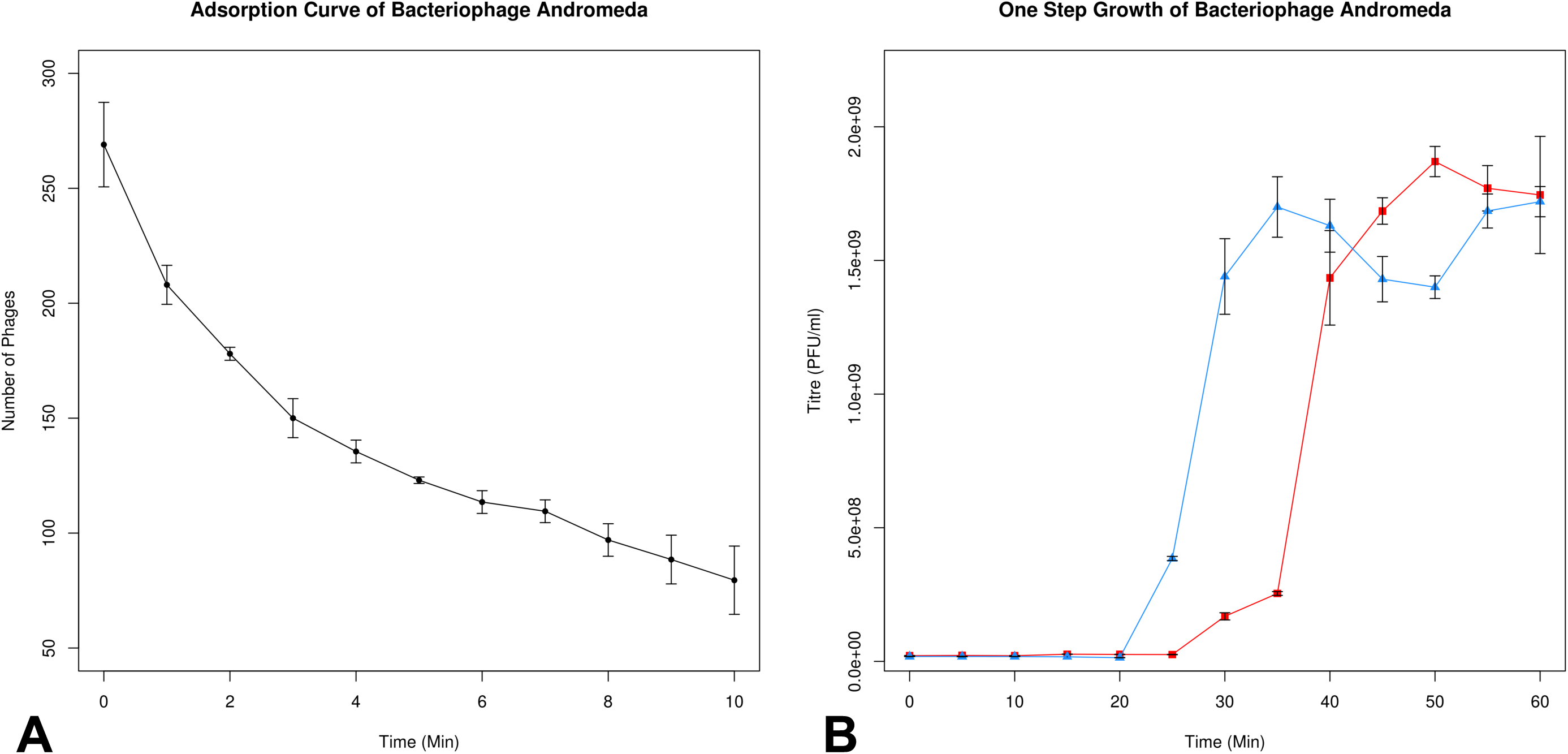
Growth characteristics of bacteriophage Andromeda. A. Determination of adsorption coefficient. B. One-step growth curves: blue line – infected *P. syringae* cells treated with chloroform (intracellular phage); red line – untreated infected *P. syringae* cells (extracellular phage). All experiments were performed in triplicates, error bars represent one standard deviation.

### Genome Structure and Gene Regulation

Phage DNA was sequenced as described in Materials and Methods to a per base average coverage of 2149. The assembly and subsequent analysis were conducted and these revealed a linear dsDNA genome of 40,008 bp (Supplementary File 1).

The GC content of the genome is 58.23% and using cumulative GC skew analysis, the putative origin of replication was found to be at 3001 bp. 46 ORFs were predicted within the Andromeda genome resulting in a total coding potential of 95.1%. Six ORFs utilise TTG, and 1 utilises GTG as start codons with the rest beginning with ATG. All ORFs are encoded off the same strand. Analysis of the Andromeda genome by both the PhageTerm [28] and Li’s methods [29] revealed that this phage is terminally redundant and likely utilises a headful mechanism of packaging similar to that of phage P1 (+ve strand analysis (p = 1.17e-4), -ve strand analysis (p = 2.81e-6)). Based on nucleotide homology, the closest relative to Andromeda is the *P. tolaasii* phage Bf7 [30] with which it shares 83% identity across 85% of its genome (Fig. 3). Of the 46 ORFs, 25 were functionally assigned and are arranged within a strict modular architecture. Eight ORFs encode products associated with DNA replication, recombination, and repair. In particular, the presence of a single subunit RNA polymerase of T3/T7 type was detected (Fig. 3; RNAP), which is located downstream of the DNA polymerase (DNAP). This RNA polymerase is conserved within *Autographivirinae* and exhibits similarity to that of phiNFS, particularly in the N- and C- termini, the specificity loop, and the palm and finger domains. Indeed, molecular modelling of both polymerases and their superimposition with TM-Align revealed remarkably similar structures (Supplementary Figure 1) with TM-score of 0.94. This, along with the characteristic position of the RNA polymerase, initially suggested the inclusion of Andromeda within the *phiKMVviruses*. Eleven ORFs in Andromeda encode structural proteins including typical gene products observed in members of the *Autographivirinae* such as tail tubular proteins. The remaining six functionally assigned ORFs encode the terminase small and large subunits as well as the components of the lysis machinery, with the lysozyme belonging to the GH24 family of proteins.

**Fig. 3.**
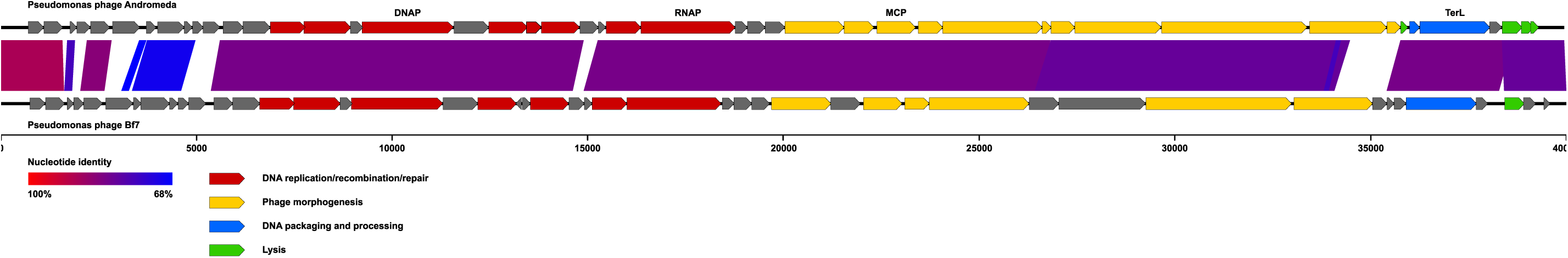
Genetic organisation of Andromeda and its homology to Pseudomonas bacteriophage Bf7. Areas of substantial similarity between two genomes are shown as trapezia connecting two genome regions coloured according to average nucleotide identity levels of these regions (as calculated by BLASTn). Arrows on the genome map represent identified genes and demonstrate the direction of their transcription. They are coloured to reflect common functions of the encoded products. Putative genes with unidentified function are shown in grey colour. DNAP – DNA polymerase, RNAP – RNA polymerase, MCP – major capsid protein, TerL – terminase large subunit.

With respect to gene regulation in Andromeda, nine σ ^70^ promoters were predicted, of which three precede the early gene region. This appears to represent a strong signal permitting the rapid transcription of early genes. Due to the presence of an RNA polymerase, we attempted to predict whether phage specific promoters were present. No statistically significant elements were found and of the obtained weak predictions, these were largely found on the non-coding strand. It is possible that Andromeda utilises σ ^70^ elements in the generation of large polycistronic mRNAs. In regard to transcriptional termination, only one rho-independent terminator was found, downstream of gp5. No tRNA genes were discovered in this phage, a feature typical of the *Autographivirinae* group of phages.

Localised single strand interruptions have been reported in *phiKMVviruses* infecting *P. aeruginosa* [2] and *P. putida* phage tf [31]. A search of the Andromeda genome failed to find the consensus associated with nicks in *P. aeruginosa* phages or *P. putida* phage. Upon subjecting Andromeda genomic DNA to NaOH denaturation, faint but distinguishable bands were observable on agarose gels which disappear when conducting this reaction after ligation (Supplementary Figure 2). These results prove that localised nicks containing adjacent 5’-phosphate and 3’-hydroxyl groups are present in the Andromeda genome. The low intensity of the bands observed may suggest that only a proportion of phage DNA molecules is nicked. This was previously demonstrated for T5 phage of *Escherichia coli* where a number of major (present in over 50% of molecules) and minor (lower presence) canonical nicks are known [32–34].

### Genetic Heterogeneity in the Andromeda Genome

It has previously been discovered that several members of the *phiKMVviruses* display a high level of genome variability that is localised to specific regions [3]. This is also true in case of Andromeda, which was found to possess the highest number of such variants. In-depth analysis of this genomic feature was conducted in order to gain insights into the nature of the changes and the potential mechanisms at play.

Analysis of the deep sequencing data of the Andromeda genome revealed that there are at least 80 positions (SNP) where nucleotide substitutions are detected with frequencies ranging from 1% to 12.48%. Detailed information with respect to each variant is provided in Supplementary Table 2. Cumulative analysis of these frequencies shows that lower frequency variants predominate, with a high level of tapering observed above 3% (Fig. 4). The mutations are distributed genome-wide, which is dissimilar to the pattern reported previously in other *phiKMVviruses* where a distinct bias of SNPs distribution (e.g. in genes responsible for phage adsorption) was observed [3]. On a cumulative basis however, it is clear that regions of high variability are observable, which suggests either the consecutive introduction of errors in a single event such as polymerase slippage, or that these represent hotspots for mutation (Fig. 4).

**Fig. 4.**
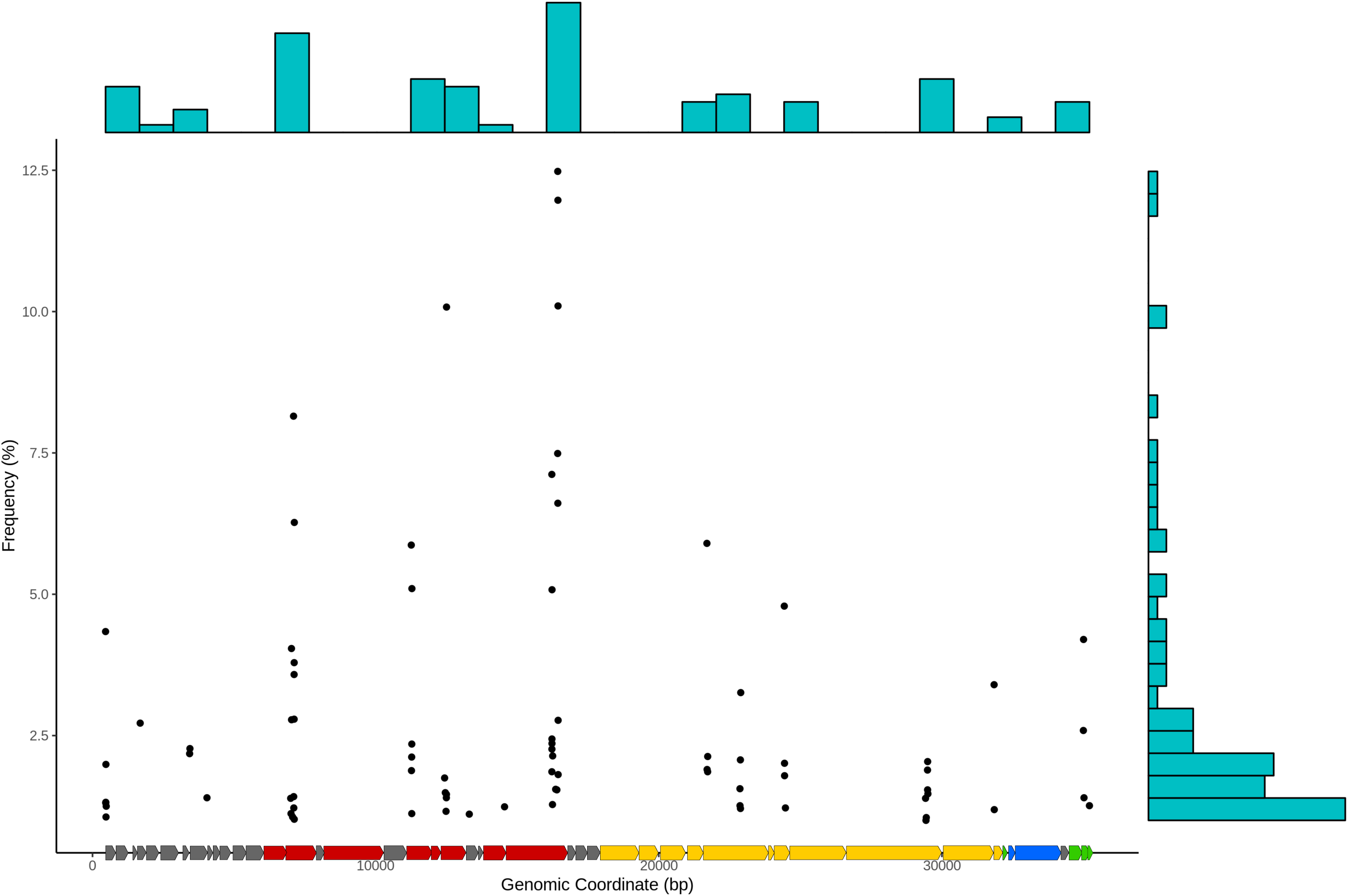
The presence and distribution of variable sites in phage Andromeda. Each dot indicates an SNP, variant frequency is given by its position along the y-axis. The histogram on the top shows uneven distribution of variable sites in the phage’s genome, the histogram on the right demonstrates the distribution of variant frequencies.

Further analysis of the nature of variants revealed that 73/80 (91.3%) lie within coding regions which is unsurprising given that the coding potential of the genome is 95.1 % (Fig. 5). With respect to those variants lying within coding regions, 55 of these result in an amino acid change. Of the 18 silent mutations, 17 were found to exist within the third codon position. This position is overall the predominant location of variants with the second codon position having a similar abundance (Fig. 5). Interestingly, these 17 silent mutations constitute only half of the total variants observed at the third position. The predominance of missense mutations in general is extremely interesting, particularly in the third position. Analysis of the proportion of transversion and transition mutations revealed an unexpected abundance of the former, with 78.75% of all variants being of this type (including those in non-coding regions). Most transversions are observed in the second codon position, closely followed by the third. In general, whilst there are more potential pathways to transversion mutations, they are in reality less common relative to transitions. The exchange of a single ring base for a double ring and vice-versa is a significant change and is more likely to result in an alteration at the amino acid level and indeed, one that is less likely to be conservative. The predominance of transversions here explains the high level of non-silent mutations observed across all codon positions and is a unique occurrence not previously reported in other biological systems. The additional abundance of transversion mutations may also further implicate the DNA polymerase as a responsible factor in variant generation through some form of biochemical bias. This is additionally supported by the presence of an insertion mutation in the Andromeda genome within ORF14 (Supplementary Table 2). This variant lies within an 8 base poly-G tract that may have given rise to a slippage event during replication and results in a +1 frameshift and truncated ORF14 product.

**Fig. 5.**
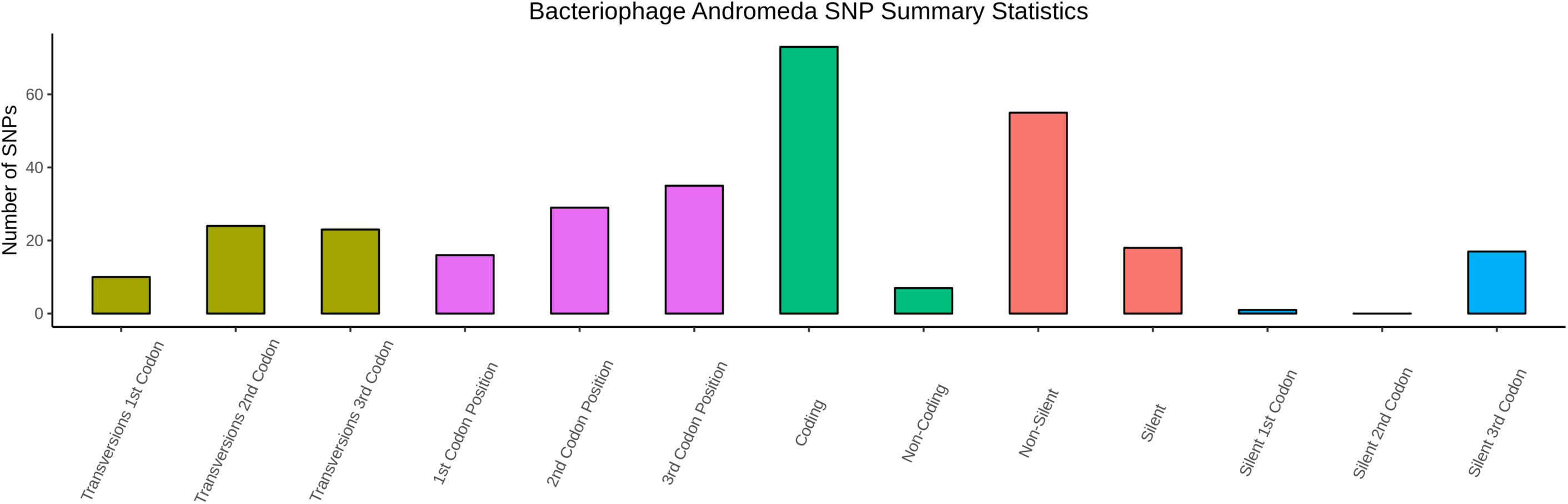
Summary statistics of SNPs in bacteriophage Andromeda with respect to their position in codons and their effect on amino acids.

The possible involvement of the phage DNA polymerases in generation of hypervariability in genomes of *phiKMVviruses* was discussed previously [3]. Phage Andromeda’s DNA polymerase is similar to those of *phiKMVviruses* with respect to absence of the thioredoxin-biding domain; however, the genome-wide distribution of variable sites in its genome suggest that different mechanisms of diversity generation might be in play in case of Andromeda.

### Phylogenetic relationships of Andromeda

*Autographivirinae* is a subfamily within *Podoviridae*, which at the time of writing included several genera, with *Teseptimaviruses, Drulisviruses, Friunaviruses, Przondoviruses, Zinderviruses* and *phiKMVviruses* being the largest. Certain features of Andromeda, such as genomic architecture and gene order, presence of nicks and hypervariable sites in its DNA, and the similarity of its gene products such as the RNA/DNA polymerases, tail tubular proteins, major capsid protein, terminase subunits, and lysis machinery, implicates this phage as a member of *Autographivirinae*, the *phiKMVviruses* in particular.

Nevertheless, it has no detectable nucleotide identity to currently recognised members of *phiKMVviruses* and has recently been assigned to *Bifseptivirus* genus within *Podoviridae* alongside with *Pseudomonas* phage Bf7 by the International Committee on Taxonomy of Viruses (ICTV proposal 2018.063B.R.Bifseptvirus, approved on 24 February 2019). This highlights the somewhat problematic nature of taxonomic classification of Andromeda and in order to better understand its phylogenetic standing, we conducted an analysis of Andromeda using two conserved proteins – RNA polymerase (RNAP) and terminase large subunit (TerL). BLASTp searches of the NCBI RefSeq database for RNAP and TerL similar to those of Andromeda (see Materials and Methods) resulted in the retrieval of 147 complete viral genomes, including 46 recognised by the ICTV (Supplementary Table 1). Almost all the phages used for phylogenetic analyses belong to *Autographivirinae* (except *Pseudomonas* phages Andromeda and Bf7 and *Aquamicrobium* phage P14) and infect a wide range of hosts from *Cyanobacteria* to *Pseudomonas*. Reconstruction of phylogenetic trees with IQ-Tree using RNAP and TerL as markers resulted in trees of similar morphology (Fig. 6a and 6b). It can be clearly seen that *Teseptimaviruses* and *Przondoviruses* cluster separately from the majority of other phages, while Andromeda belongs to a loose cluster of *Autographivirinae* phages comprising several smaller genera with only few members (e.g. *Bifseptivirus* – *Pseudomonas* phages Andromeda and Bf7, *Aqualcavirus* – *Aquamicrobium* phage P14) and unclassified *Autographivirinae* (e.g. *Ralstonia* phage RSB3). Although we do not intend to propose any novel classification, we designate this cluster of viruses as “ExophiKMVviruses” for the purposes of discussing this group and to highlight its distinctive position relative to other *Autographivirnae*.

**Fig. 6.**
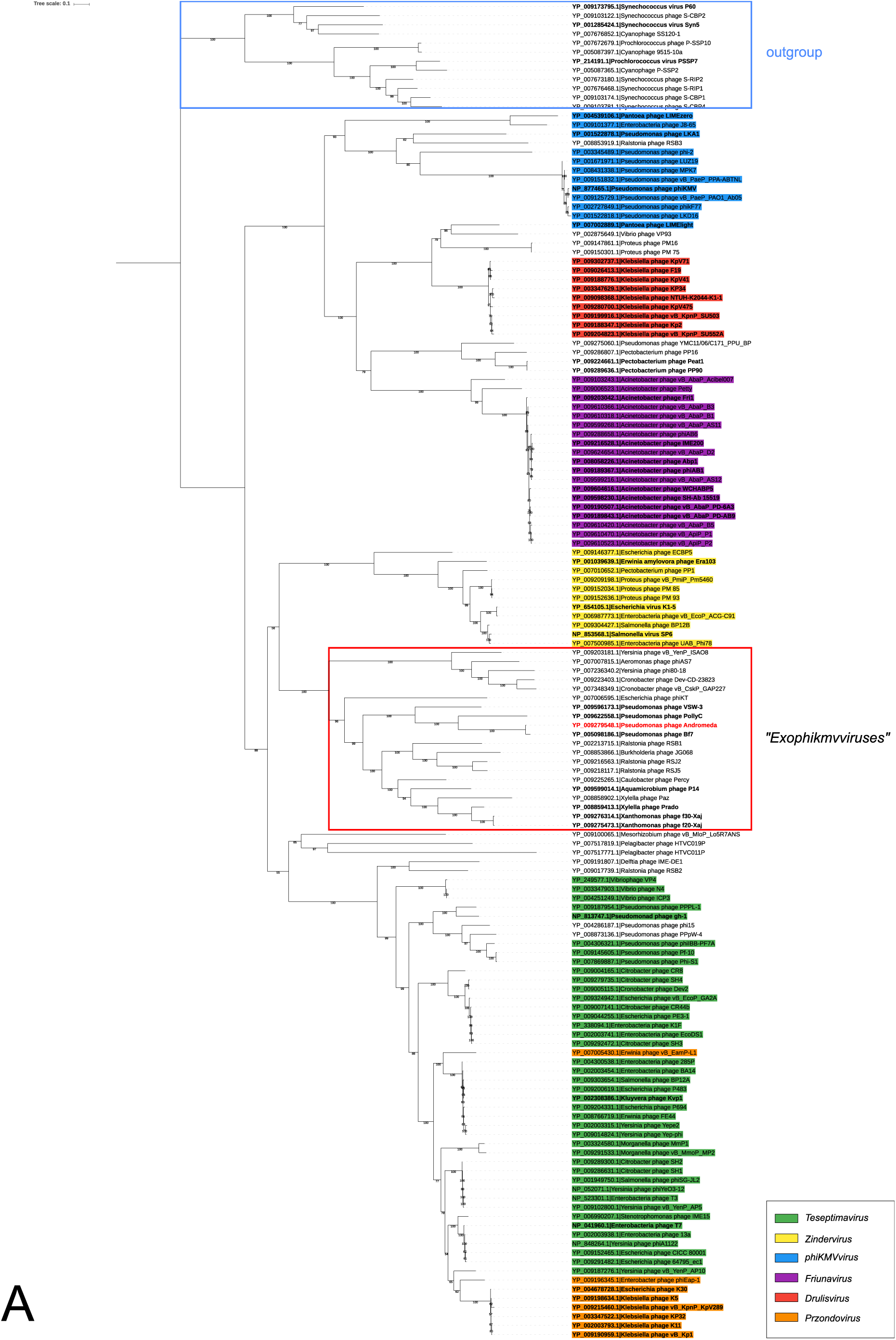
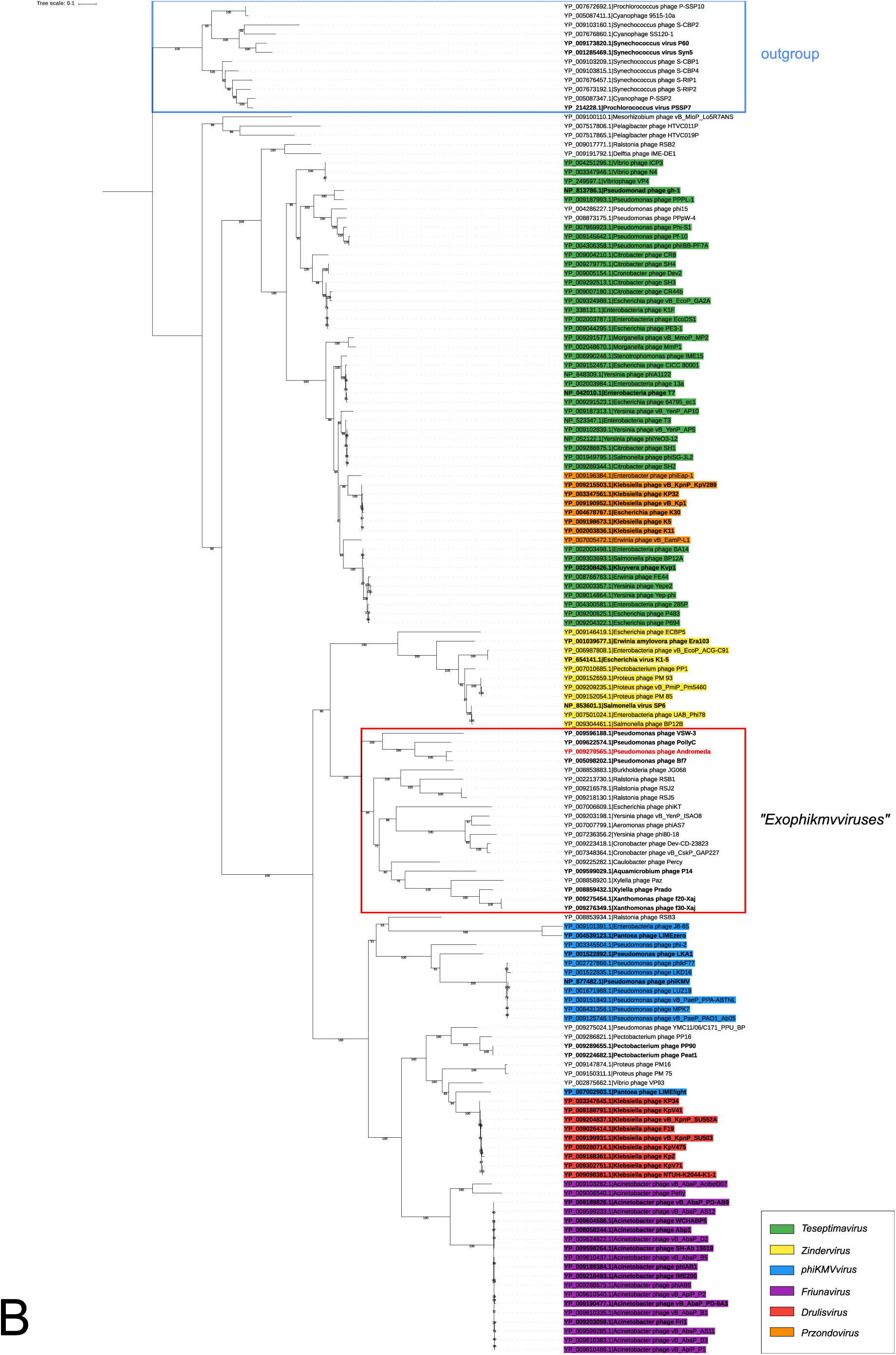
Analysis of phylogenetic relationships of phage Andromeda. A. Consensus phylogenetic tree based on multiple sequence alignment of RNAP amino acid sequences of 147 RefSeq phages; LG+F+R6 substitution model. B. Consensus phylogenetic tree based on multiple sequence alignment of TerL amino acid sequences of 147 RefSeq phages; LG+R5 substitution model. Consensus trees were constructed from 1000 bootstrap trees, numbers in parentheses are bootstrap supports (%). Both trees were rooted with a cluster of 11 cyanophages as outgroup. “Exophikmvviruses” are outlined by a red box.

Despite not currently being classified by ICTV as *Autographivirinae*, phage Andromeda as well as Bf7 could be included as members of this group based on the presence of a single subunit RNAP in their genomes and their position within the cluster of “ExophiKMVviruses”. When we analyse the gene order of representatives of each genus (Fig. 7), we find that the “ExophiKMVviruses”, despite being most closely related to *Zinderviruses* based on sequence identity percent of marker genes, show the downstream localisation of the single-subunit RNA polymerase as is typical for “true” *phiKMVviruses*. This is a feature conserved across the group (Fig. 7). Analysis of genome size distribution of the putative *Autographivirinae* (Fig. 8) show that “ExophiKMVviruses” possess a genome size range similar to that of other *phiKMVviruses*. We have shown previously that *phiKMVviruses* such as *Pseudomonas* phage phiNFS lack the thioredoxin binding domain in their DNA polymerases [3] that is responsible for heightened processivity and fidelity of *Teseptimaviruses* [35, 36]. Molecular modelling and superimposition of the Andromeda and T7 DNA polymerases shows the absence of this domain in Andromeda and highlights another difference between *phiKMVviruses* and “Exophikmvviruses” in comparison to *Teseptimaviruses* (Supplementary Figure 3). Taken together, it seems that the “ExophiKMVviruses” might occupy an interface group between the *phiKMVviruses* and the *Teseptimaviruses*, although sequence based classification alone being insufficient to adequately resolve the relationships of the *Autographivirinae* as a whole.

**Fig. 7.**
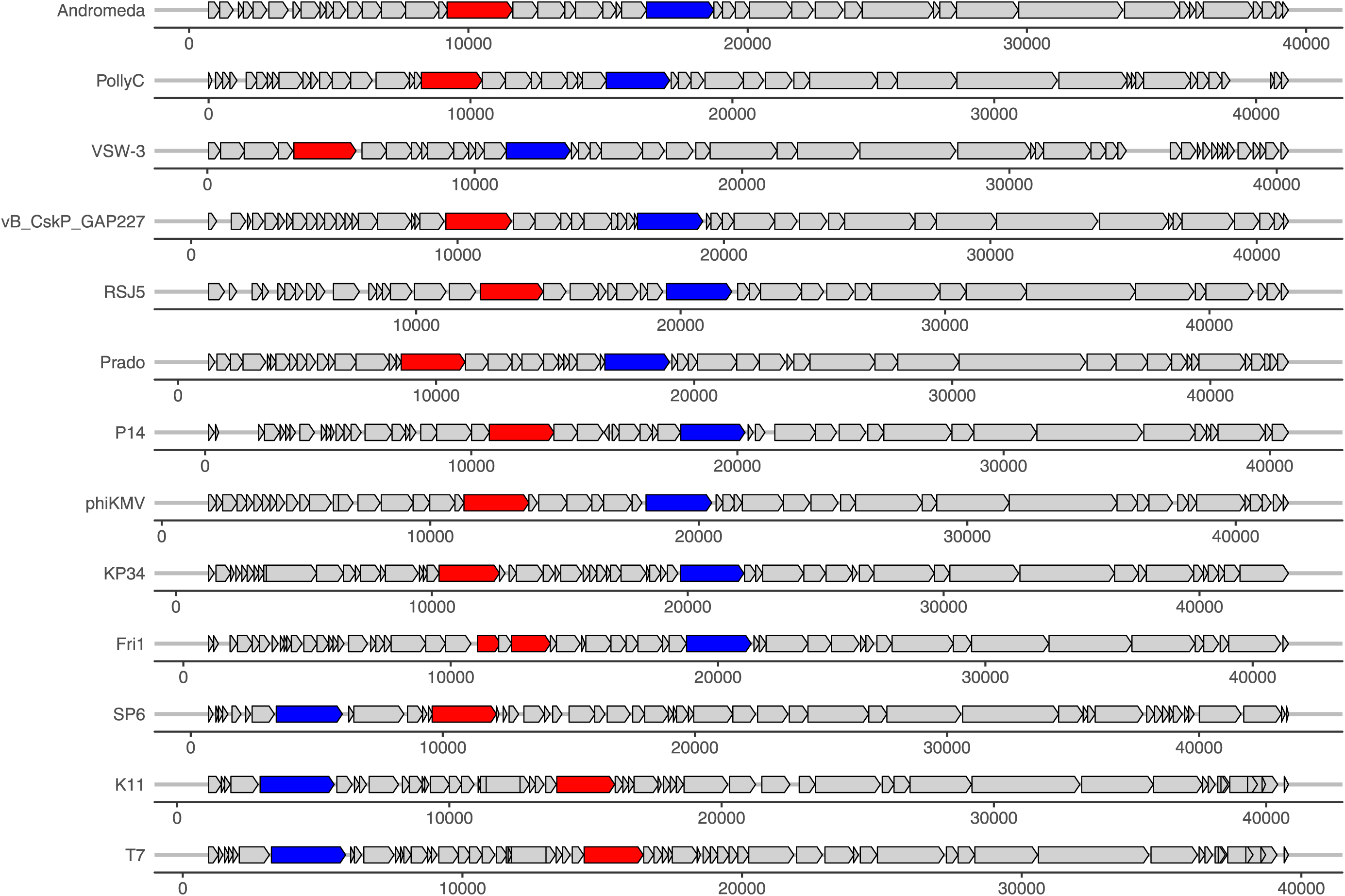
Comparison of positions of RNA (blue) and DNA (red) polymerases in representatives of different genera. In *Zinderviruses* (SP6), *Teseptimaviruses* (T7) and *Przondoviruses* (K11) DNA polymerase is adjacent to structural genes and is positioned downstream of RNA polymerase, while for the rest of the viruses the opposite is true.

**Fig. 8.**
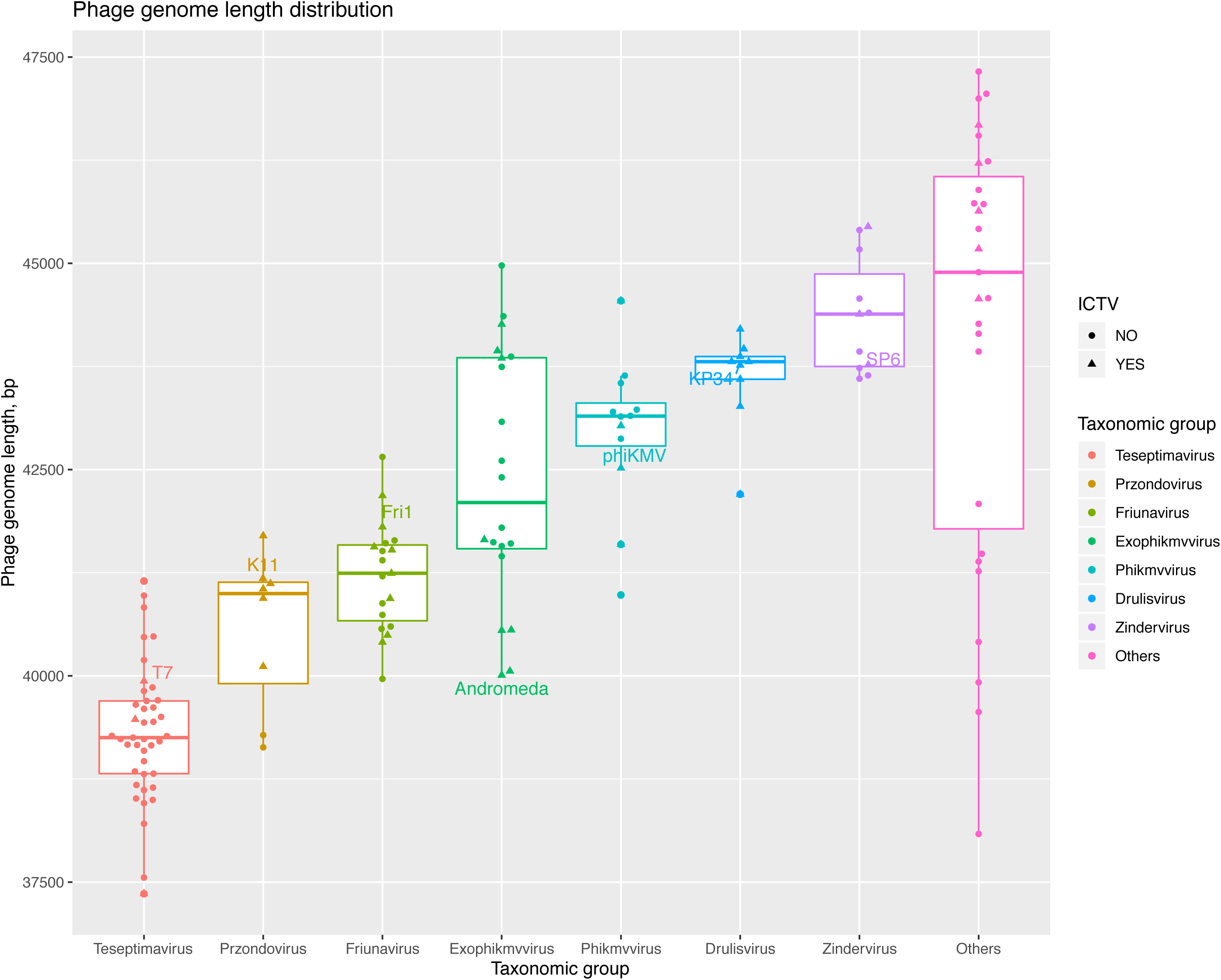
Genome size distribution among viruses of different taxonomic groups analysed in this study.

Our analysis of the Andromeda phage has once again highlighted the existence of a novel phenomenon within *Pseudomonas* phages that results in the production of highly heterogeneous populations. Whilst error-prone polymerase activity is almost certainly to be involved in the generation of genomic hypervariability, it unlikely to be the only constituent at play, with intriguing patterns of codon position distribution and localisation suggesting the involvement of some post-replication mechanisms. The taxonomic position of Andromeda suggests that this phenomenon is not unique to *phiKMVviruses* and could potentially include “Exophikmvviruses” and/or *Autographivirinae* as a whole.

## Authors and contributors

Conceptualisation: D.J.M. and L.A.K.; data collection and analysis: D.J.M. and T.A.S.; writing – original draft preparation: D.J.M., T.A.S. and L.A.K.; writing – review and editing: T.A.S., D.J.M. and L.A.K

## Funding information

This work received no specific grant from any funding agency

## Conflicts of interest

The authors declare that there are no conflicts of interest.

## Ethical statement

No experimental work with humans or animals was performed in this study.

## Legends to Supplementary Data

**Supplementary Figure 1**. Superimposition of three-dimensional structures of RNA polymerases from Pseudomonas phages Andromeda (salmon) and phiNFS (purple).

**Supplementary Figure 2**. Visualisation of the localised single stranded interruptions in the Andromeda’s genome. (A) GeneRuler 1 kb Plus DNA Ladder ladder; (B) Untreated phage DNA; (C) NaOH treated DNA, major DNA hydrolysis products are labelled (1-3); (D) NaOH treated DNA which was preliminarily ligated by T4 DNA ligase.

**Supplementary Figure 3**. Superimposition of three-dimensional structures of DNA polymerases from Pseudomonas phage Andromeda (salmon) and Enterobacteria phage T7 (blue, model derived from PDB 1SKR).

**Supplementary Table 1**. NCBI RefSeq phage genomes used in this study.

**Supplementary Table 2**. Detailed description of SNPs and indels in phage Andromeda genome.

**Supplementary File 1**. Phage Andromeda annotated genome (Genbank format).

## Notes

#### Summary of Updates

List of authors updated (change of corresponding author)

## References

1. Kulakov LA, Ksenzenko VN, Kochetkov VV, Mazepa VN, Boronin AM. DNA homology and adsorption specificity of *Pseudomonas aeruginosa* virulent bacteriophages. MGG Mol Gen Genet 1985; 200: 123–127.

2. Kulakov LA, Ksenzenko VN, Shlyapnikov MG, Kochetkov VV, Del Casale A, Allen CCR, et al. Genomes of “phiKMV-like viruses” of *Pseudomonas aeruginosa* contain localized single-strand interruptions. Virology 2009; 391: 1–4.

3. Magill DJ, Kucher PA, Krylov VN, Pleteneva EA, Quinn JP, Kulakov LA. Localised genetic heterogeneity provides a novel mode of evolution in dsDNA phages. Sci Rep 2017; 7.

4. Sambrook Joseph, Russell DW. Molecular cloning□: a laboratory manual. 2001. Cold Spring Harbor Laboratory Press, Cold Spring Harbor, N.Y.

5. Clokie MRJ, Kropinski AM (eds). Bacteriophages: methods and protocols. Volume 1. 2009. Humana Press, New York.

6. Bolger AM, Lohse M, Usadel B. Trimmomatic: a flexible trimmer for Illumina sequence data. Bioinformatics 2014; 30: 2114–2120.

7. Bankevich A, Nurk S, Antipov D, Gurevich AA, Dvorkin M, Kulikov AS, et al. SPAdes: A New Genome Assembly Algorithm and Its Applications to Single-Cell Sequencing. J Comput Biol 2012; 19: 455–477.

8. Boetzer M, Henkel CV, Jansen HJ, Butler D, Pirovano W. Scaffolding pre-assembled contigs using SSPACE. Bioinforma Oxf Engl 2011; 27: 578–579.

9. Thorvaldsdóttir H, Robinson JT, Mesirov JP. Integrative Genomics Viewer (IGV): high-performance genomics data visualization and exploration. Brief Bioinform 2013; 14: 178–192.

10. Carver T, Harris SR, Berriman M, Parkhill J, McQuillan JA. Artemis: an integrated platform for visualization and analysis of high-throughput sequence-based experimental data. Bioinformatics 2012; 28: 464–469.

11. Lukashin AV, Borodovsky M. GeneMark.hmm: New solutions for gene finding. Nucleic Acids Res 1998; 26: 1107–1115.

12. Altschul SF, Gish W, Miller W, Myers EW, Lipman DJ. Basic local alignment search tool. J Mol Biol 1990; 215: 403–410.

13. Boratyn GM, Schäffer AA, Agarwala R, Altschul SF, Lipman DJ, Madden TL. Domain enhanced lookup time accelerated BLAST. Biol Direct 2012; 7: 12–12.

14. Jones P, Binns D, Chang H-Y, Fraser M, Li W, McAnulla C, et al. InterProScan 5: genome-scale protein function classification. Bioinforma Oxf Engl 2014; 30: 1236–1240.

15. Zimmermann L, Stephens A, Nam S-Z, Rau D, Kübler J, Lozajic M, et al. A Completely Reimplemented MPI Bioinformatics Toolkit with a New HHpred Server at its Core. J Mol Biol 2018; 430: 2237–2243.

16. Lowe TM, Eddy SR. tRNAscan-SE: a program for improved detection of transfer RNA genes in genomic sequence. Nucleic Acids Res 1997; 25: 955–964.

17. Laslett D, Canback B. ARAGORN, a program to detect tRNA genes and tmRNA genes in nucleotide sequences. Nucleic Acids Res 2004; 32: 11–16.

18. Naville M, Ghuillot-Gaudeffroy A, Marchais A, Gautheret D. ARNold: A web tool for the prediction of Rho-independent transcription terminators. RNA Biol 2011; 8: 11–13.

19. Zuker M. Mfold web server for nucleic acid folding and hybridization prediction. Nucleic Acids Res 2003; 31: 3406–3415.

20. Bailey TL, Johnson J, Grant CE, Noble WS. The MEME Suite. Nucleic Acids Res 2015; 43: W39–W49.

21. Katoh K, Standley DM. MAFFT Multiple Sequence Alignment Software Version 7: Improvements in Performance and Usability. Mol Biol Evol 2013; 30: 772–780.

22. Capella-Gutiérrez S, Silla-Martínez JM, Gabaldón T. trimAl: a tool for automated alignment trimming in large-scale phylogenetic analyses. Bioinformatics 2009; 25: 1972–1973.

23. Nguyen L-T, Schmidt HA, von Haeseler A, Minh BQ. IQ-TREE: A Fast and Effective Stochastic Algorithm for Estimating Maximum-Likelihood Phylogenies. Mol Biol Evol 2015; 32: 268–274.

24. Letunic I, Bork P. Interactive Tree Of Life (iTOL) v4: recent updates and new developments. Nucleic Acids Res 2019; 47: W256–W259.

25. Yang J, Yan R, Roy A, Xu D, Poisson J, Zhang Y. The I-TASSER Suite: protein structure and function prediction. Nat Methods 2015; 12: 7–8.

26. Pettersen EF, Goddard TD, Huang CC, Couch GS, Greenblatt DM, Meng EC, et al. UCSF Chimera—A visualization system for exploratory research and analysis. J Comput Chem 2004; 25: 1605–1612.

27. Zhang Y, Skolnick J. TM-align: a protein structure alignment algorithm based on the TM-score. Nucleic Acids Res 2005; 33: 2302–2309.

28. Garneau JR, Depardieu F, Fortier L-C, Bikard D, Monot M. PhageTerm: a tool for fast and accurate determination of phage termini and packaging mechanism using next-generation sequencing data. Sci Rep 2017; 7: 8292.

29. Li S, Fan H, An X, Fan H, Jiang H, Chen Y, et al. Scrutinizing Virus Genome Termini by High-Throughput Sequencing. PLOS ONE 2014; 9: e85806.

30. Sajben-Nagy E, Maróti G, Kredics L, Horváth B, Párducz Á, Vágvölgyi C, et al. Isolation of new Pseudomonas tolaasii bacteriophages and genomic investigation of the lytic phage BF7. FEMS Microbiol Lett 2012; 332: 162–169.

31. Glukhov AS, Krutilina AI, Shlyapnikov MG, Severinov K, Lavysh D, Kochetkov VV, et al. Genomic Analysis of Pseudomonas putida Phage tf with Localized Single-Strand DNA Interruptions. PLOS ONE 2012; 7: e51163.

32. Bujard H. LOCATION OF SINGLE-STRAND INTERRUPTIONS IN THE DNA OF BACTERIOPHAGE T5+. Proc Natl Acad Sci U S A 1969; 62: 1167–1174.

33. Hayward GS, Smith MG. The chromosome of bacteriophage T5: I. Analysis of the single-stranded DNA fragments by agarose gel electrophoresis. J Mol Biol 1972; 63: 383–395.

34. Johnston JV, Nichols BP, Donelson JE. Distribution of ‘minor’ nicks in bacteriophage T5 DNA. J Virol 1977; 22: 510–519.

35. Magill DJ, McGrath JW, O’Flaherty V, Quinn JP, Kulakov LA. Insights into the structural dynamics of the bacteriophage T7 DNA polymerase and its complexes. J Mol Model 2018; 24: 144.

36. Tabor S, Huber HE, Richardson CC. Escherichia coli thioredoxin confers processivity on the DNA polymerase activity of the gene 5 protein of bacteriophage T7. J Biol Chem 1987; 262: 16212–16223.

