## Supplementary Figure 1 for "Genomic hypervariability of phage Andromeda is unique among known dsDNA viruses"

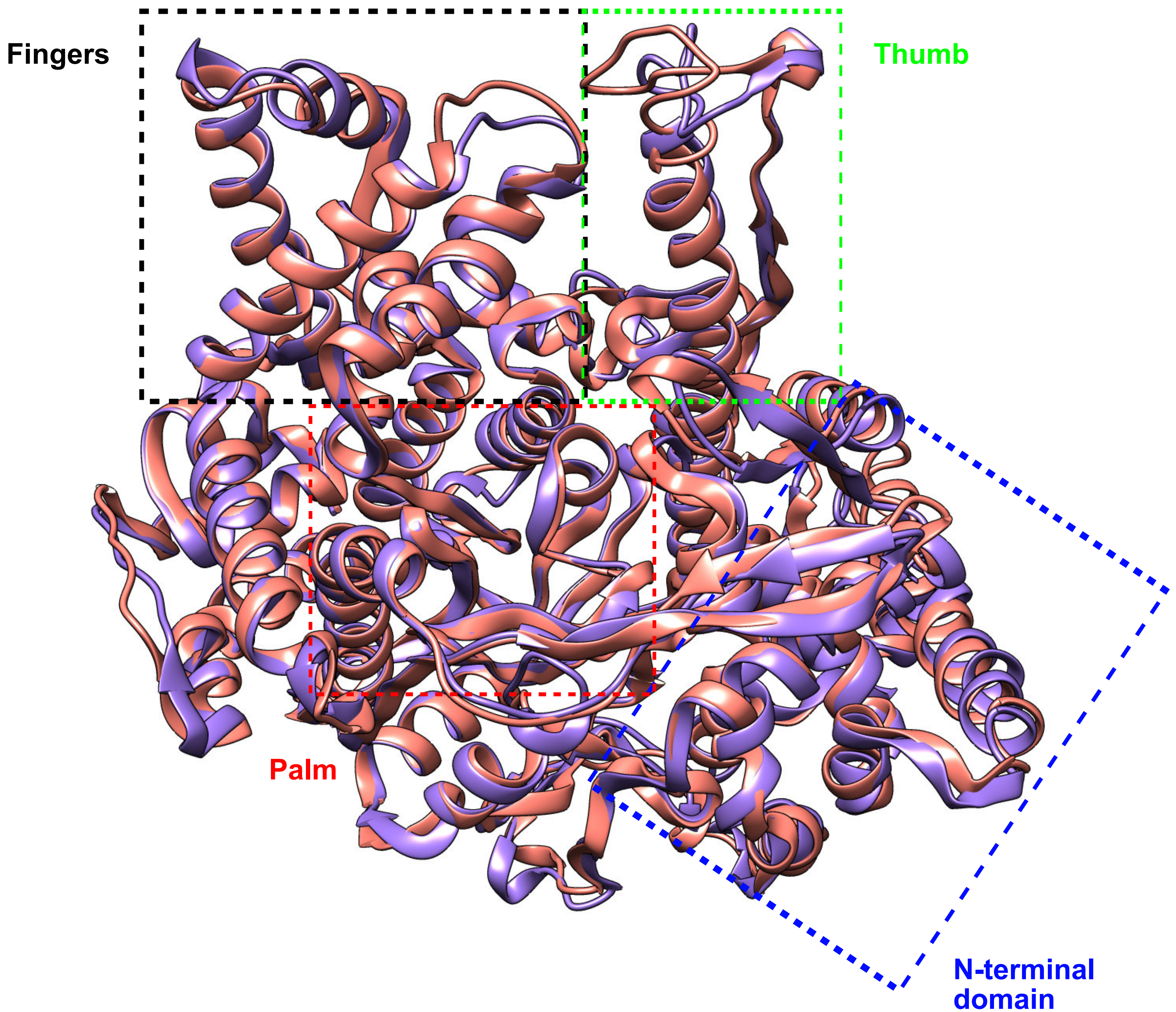


Andromeda RNA polymerase

phiNFS RNA polymerase

```
*********************************************************************
*                        TM-align (Version 20190822)                 *
* An algorithm for protein structure alignment and comparison        *
* Based on statistics:                                               *
*       0.0 < TM-score < 0.30, random structural similarity          *
*       0.5 < TM-score < 1.00, in about the same fold                *
* Reference: Y Zhang and J Skolnick, Nucl Acids Res 33, 2302-9 (2005)*
* Please email your comments and suggestions to:      *
*********************************************************************

Name of Chain_1: Andromeda RNA polymerase
Name of Chain_2: phiNFS RNA polymerase
Length of Chain_1:  803 residues
Length of Chain_2:  816 residues

Aligned length=  786, RMSD=   1.70, Seq_ID=n_identical/n_aligned= 0.265
TM-score= 0.95747 (if normalized by length of Chain_1)
TM-score= 0.94243 (if normalized by length of Chain_2)
(You should use TM-score normalized by length of the reference protein)
```
