## Supplementary figures and images for "Genomic hypervariability of phage Andromeda is unique among known dsDNA viruses"

### Supplementary Figure 2

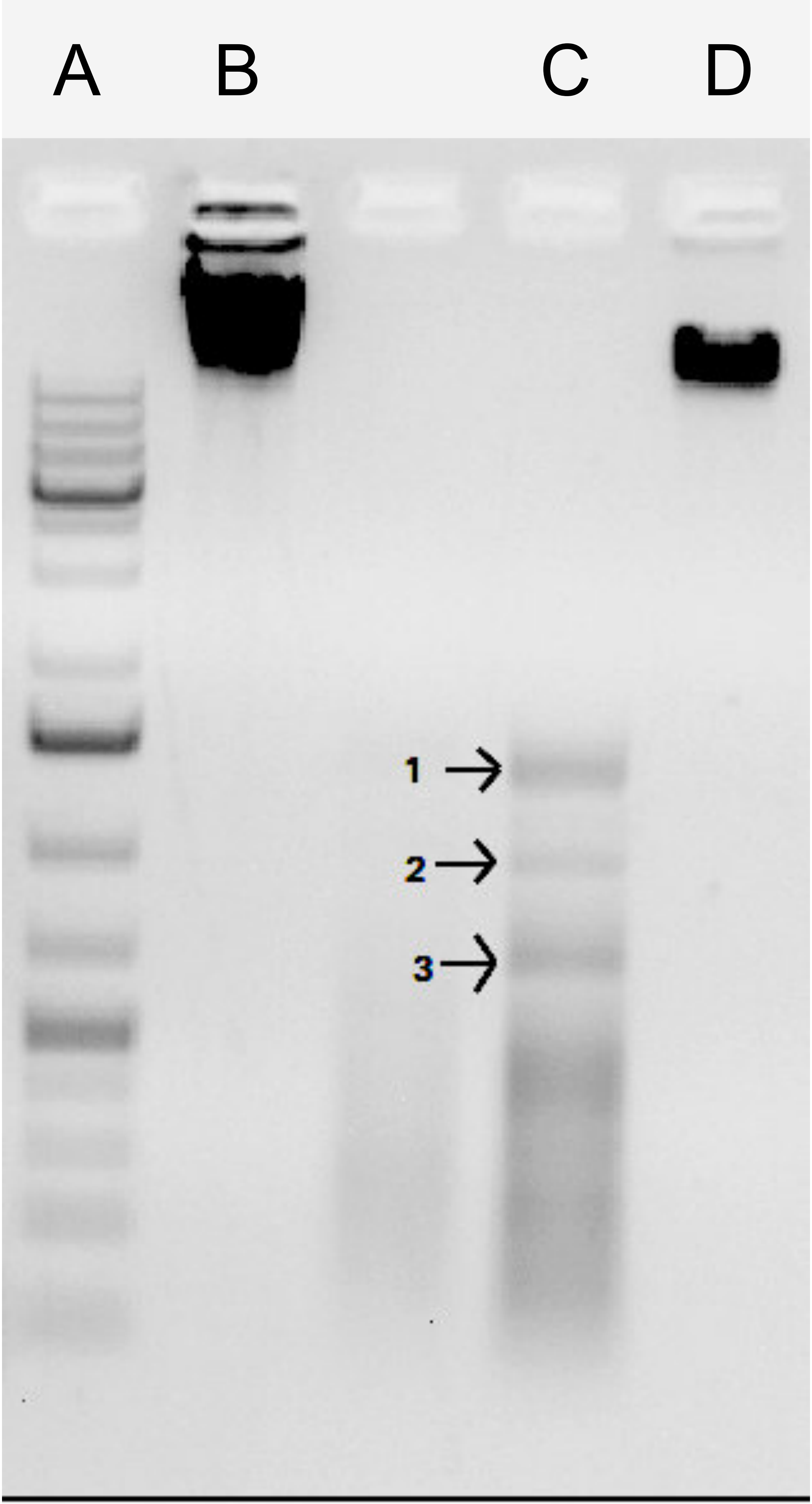
