## Supplementary Figure 3 for "Genomic hypervariability of phage Andromeda is unique among known dsDNA viruses"

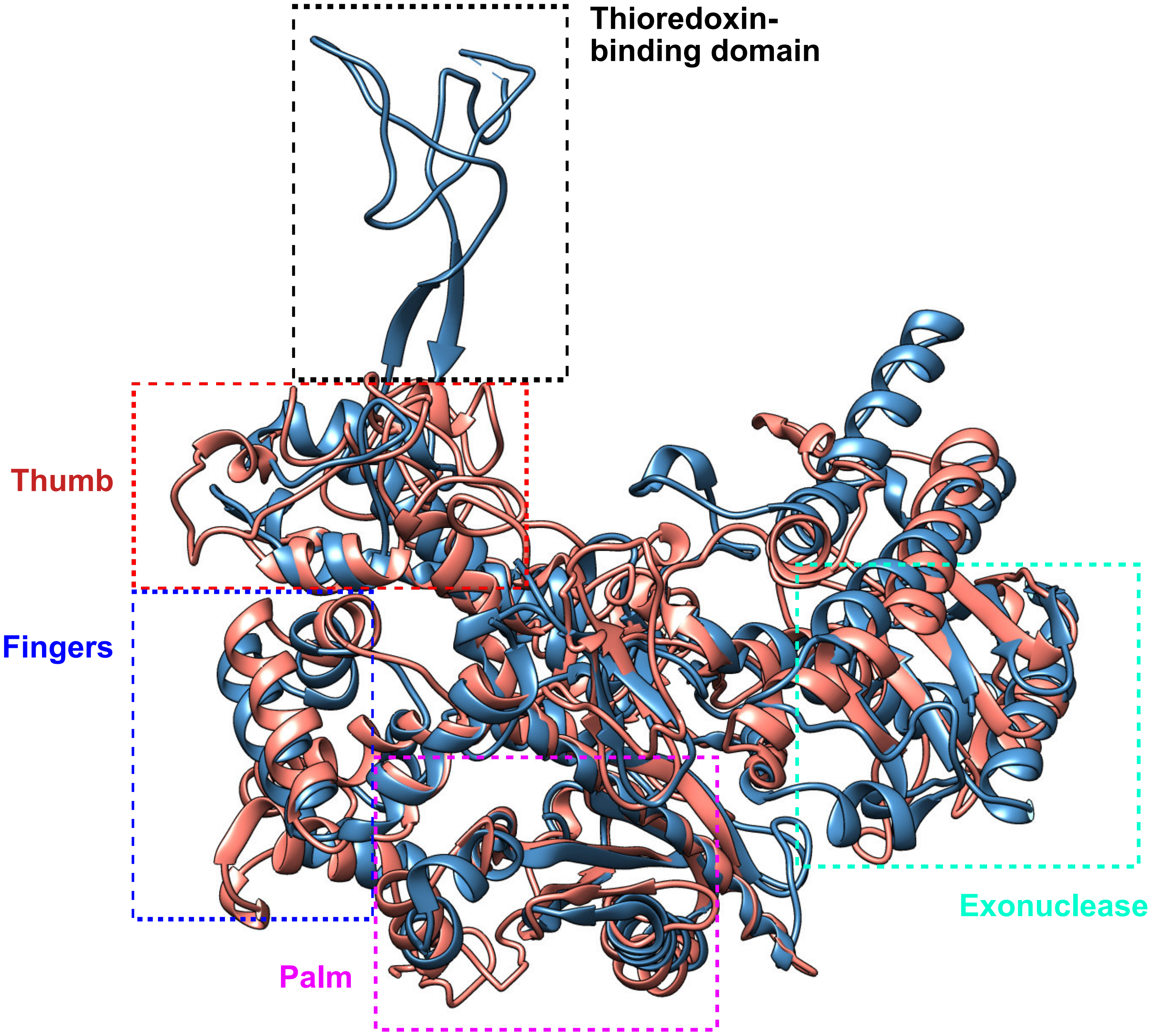


Andromeda DNA polymerase

T7 DNA polymerase

```
 **************************************************************************
 *                        TM-align (Version 20190822)                     *
 * An algorithm for protein structure alignment and comparison            *
 * Based on statistics:                                                   *
 *       0.0 < TM-score < 0.30, random structural similarity              *
 *       0.5 < TM-score < 1.00, in about the same fold                    *
 * Reference: Y Zhang and J Skolnick, Nucl Acids Res 33, 2302-9 (2005)    *
 * Please email your comments and suggestions to:          *
 **************************************************************************

Name of Chain_1: Andromeda DNA polymerase
Name of Chain_2: T7 DNA polymerase
Length of Chain_1:  780 residues
Length of Chain_2:  669 residues

Aligned length=  568, RMSD=   4.10, Seq_ID=n_identical/n_aligned= 0.164
TM-score= 0.64191 (if normalized by length of Chain_1)
TM-score= 0.73925 (if normalized by length of Chain_2)
(You should use TM-score normalized by length of the reference protein)
```
